# A Ribosomal RNA modification regulates translation of PI3K/AKT activators in high-grade serous ovarian carcinoma

**DOI:** 10.64898/2026.10.06.757003

**Authors:** Andrea Rivera-Pizana, Laurence Faucher-Giguère, Michelle S. Scott, Sherif Abou Elela

## Abstract

PI3K/AKT signalling is a key driver of ovarian cancer progression and an important therapeutic target. Its deregulation is usually explained by genetic changes and altered signalling at the cell surface. Here, we identify an additional layer of pathway control, in which a change in the level of a cancer-associated ribosomal RNA modification alters the translation of proteins that regulate PI3K/AKT activity. SNORA81 is a high-grade serous ovarian carcinoma-associated H/ACA small nucleolar RNA that directs pseudouridylation of 28S ribosomal RNA at position U4636. Integration of tumour transcriptomes with independent SNORA81-depletion datasets identified 22 SNORA81-linked transcripts with reversed disease-associated expression, 19 of them linked to PI3K/AKT signalling. SNORA81 depletion reduced AKT phosphorylation, impaired cell growth, migration, and altered cell-cycle distribution. Molecularly, polysome profiling following SNORA81 depletion identified 524 mRNAs with altered ribosome association, 485 of which changed independently of RNA abundance and included 29 PI3K/AKT regulators. SNORA81 depletion reduced ribosome association of ITGB4, RASGRP2, IGF1 and FGF18 transcripts, and reduced ITGB4 and RASGRP2 protein abundance. IRAK1 and SPC24 acted as downstream RNA-response genes contributing to selected SNORA81-associated phenotypes. ITGB4 expression partially restored cell growth and restored downstream RNA responses after SNORA81 depletion. These findings identify rRNA modification as a translational regulator of PI3K/AKT signalling in ovarian cancer.

## INTRODUCTION

Ovarian cancer remains one of the most lethal gynaecological malignancies, with high-grade serous ovarian carcinoma (HGSC) as its most common subtype that accounts for most ovarian cancer deaths [1]. HGSC is frequently diagnosed at an advanced stage and is characterised by extensive genomic instability, rapid dissemination and frequent relapse after treatment [1, 2]. Its molecular landscape includes near-universal *TP53* mutation [3], homologous-recombination deficiency in approximately half of tumours and widespread somatic copy-number alterations [4]. However, these genomic features do not fully explain the signalling activities that underlie the aggressive behaviour of HGSC.

The phosphoinositide 3-kinase (PI3K)/AKT pathway is frequently altered in ovarian cancer and remains an important therapeutic target, although pathway inhibitors have shown limited benefit in unselected patients [5]. In the canonical pathway, growth factors activate receptor tyrosine kinases, whereas integrins signal through adhesion-associated proteins such as FAK and SRC. These signals, together with RAS-dependent inputs, converge on PI3K, which converts PIP2 into PIP3. PIP3 recruits AKT to the plasma membrane, where phosphorylation by PDK1 and mTORC2 activates downstream programmes controlling tumour-cell growth, survival and invasion. PI3K/AKT activity in ovarian cancer is commonly attributed to genomic alterations in pathway components, including copy-number gain of *PIK3CA* and related subunits [6], and to loss of PTEN-mediated restraint, detected in over a third of high-grade serous tumours [7]. Nevertheless, pathway activity varies across ovarian tumours, defines prognostically distinct subgroups and is not always predicted by canonical genomic alterations [8]. Whether translation of upstream signalling proteins provides an additional level of PI3K/AKT regulation remains unclear.

Protein abundance is determined not only by transcription and protein turnover but also by mRNA translation [9]. Cancer cells reprogram translation to sustain the production of selected proteins under stress, supporting migration, invasion, metastasis and adaptation to the tumour microenvironment [9, 10]. Ribosomes are also increasingly recognised as heterogeneous molecular machines whose composition, associated factors and ribosomal RNA (rRNA) modifications may influence the translation of particular mRNAs [11, 12]. Moreover, rRNA modifications are not uniform: many sites are only modified in a subset of ribosomes and their levels vary with developmental and disease context [13, 14]. Such sites are referred to as partially modified sites. Variation in modification stoichiometry could therefore generate functionally distinct ribosome states and alter signalling-protein output without producing a global change in protein synthesis. However, whether cancer-associated rRNA modifications regulate the translation of proteins controlling specific oncogenic pathways remains poorly understood.

Small nucleolar RNAs (snoRNAs) guide rRNA maturation and modification: box C/D snoRNAs direct 2′-O-methylation, whereas H/ACA snoRNAs direct pseudouridylation [14]. Although altered snoRNA expression and rRNA modification are increasingly associated with cancer [12, 15], the mechanisms connecting individual modifications to oncogenic signalling remain largely unresolved. We previously identified SNORA81 as an HGSC-associated H/ACA snoRNA that directs pseudouridylation of 28S rRNA U4636 (U4606 in the rRNA annotation used in our previous study) [16]. SNORA81 expression distinguishes HGSC from serous borderline tumours (SBT) and normal fallopian tube epithelium, and SNORA81 depletion reduces U4636 pseudouridylation, proliferation and migration in ovarian cancer cells [16]. These findings establish a connection between SNORA81-directed rRNA modification and HGSC-associated phenotypes but do not explain the underlying signalling mechanism (**Figure 1A**).

**Figure 1.**
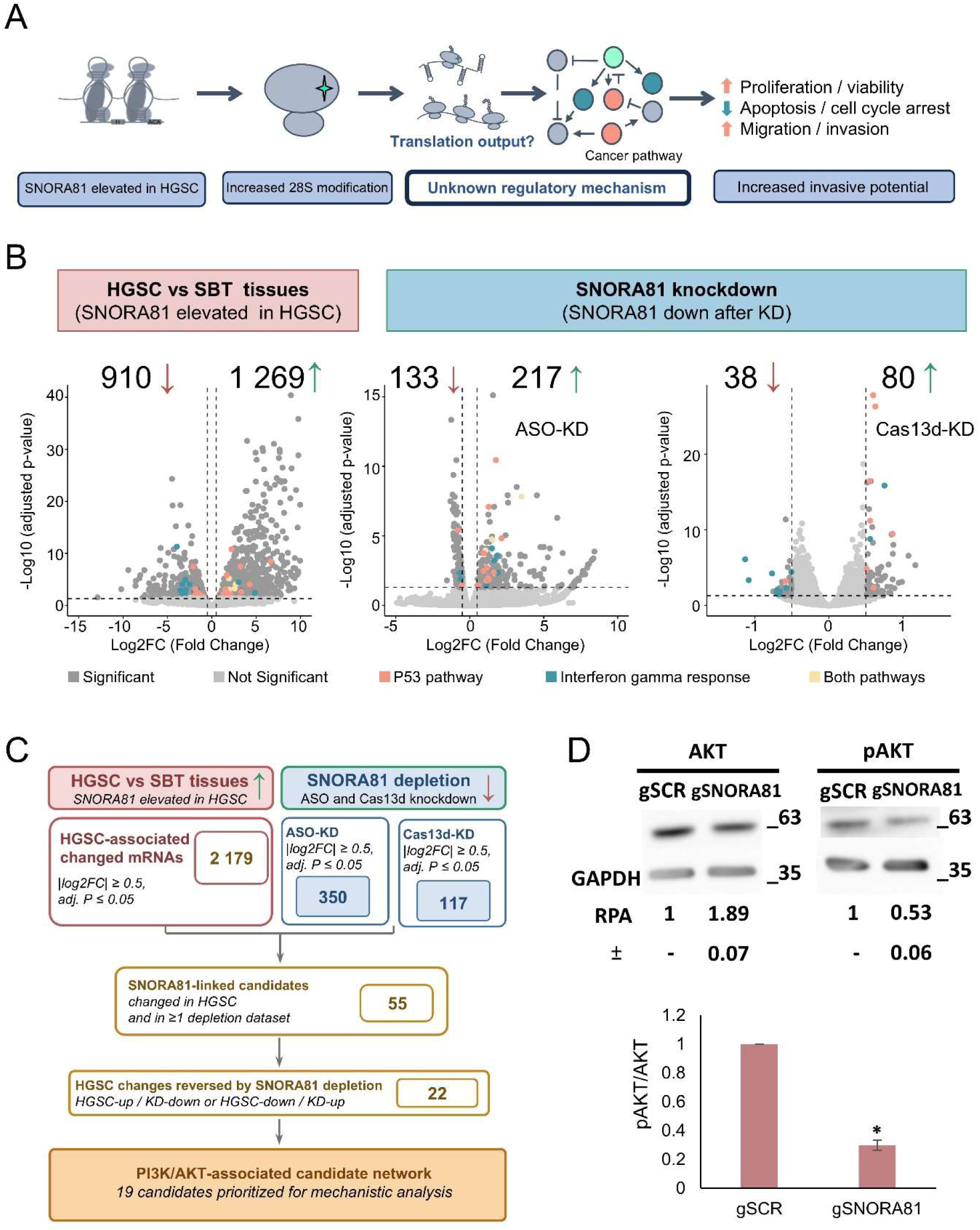
SNORA81 depletion remodels PI3K/AKT-associated gene expression in HGSC. **A**, Working model and mechanistic gap addressed in this study. Increased SNORA81 in HGSC directs pseudouridylation of 28S rRNA at position U4636 and is associated with increased proliferation and viability, reduced apoptosis and increased migration and invasion [16]; the intervening regulatory mechanism is unknown. Arrows, activation; blunt-ended lines, inhibition. **B**, Differential mRNA abundance in tumour tissues and after SNORA81 depletion. Volcano plots for SBT relative to HGSC tissues (left) and for ASO-(middle) or Cas13d-mediated (right) SNORA81 knockdown (KD) relative to matched controls in OVCAR-3 cells. Significance thresholds were |log2 fold change| ≥ 0.5 and adjusted *P* ≤ 0.05. Positive log2 fold change denotes higher abundance in SBT or in depleted cells. In SBT versus HGSC, 1 269 mRNAs were increased and 910 decreased; after ASO depletion, 217 were increased and 133 decreased; after Cas13d depletion, 80 were increased and 37 decreased. Colours denote annotation to the p53 pathway (pink), the interferon-γ response (blue), or both (yellow). **C**, Workflow integrating the three datasets: 55 mRNAs were altered in the tissue comparison and in at least one depletion dataset, 22 showed opposite directionality between HGSC and depleted cells, and of those, 19 mapped to PI3K/AKT-associated signalling. **D**, AKT phosphorylation after SNORA81 depletion. Western blot of total AKT, phospho-AKT (Thr308) and GAPDH in OVCAR-3 cells transfected with Cas13d and gSNORA81 or scrambled control (gSCR). Phospho-AKT and total AKT were detected on separate blots from the same experiment, processed in parallel. Relative protein abundance, normalised to GAPDH and to gSCR, is shown beneath each blot. Bar graph shows the phospho-AKT/AKT ratio; mean ± s.d., n = 3 biologically independent experiments; two-tailed unpaired *t*-test, *P* = 0.02. Blots were cropped for clarity.

Here, we asked whether SNORA81-dependent rRNA modification regulates PI3K/AKT signalling by altering translation of upstream pathway activators. We integrated tumour transcriptomes with two independent SNORA81-depletion datasets to identify disease-associated RNA responses, measured AKT phosphorylation as a pathway-activity readout and tested selected downstream response genes. We then combined matched RNA sequencing, polysome profiling and protein analyses to distinguish changes in RNA abundance from translational regulation. Our results identify a translational layer in which the SNORA81-associated ribosome state regulates upstream PI3K/AKT activators and connects selective translation to downstream RNA-response programmes and HGSC-associated cellular phenotypes.

## MATERIALS AND METHODS

### Tissue sourcing and ethics

HGSC and SBT tissues were provided by the Ontario Tumour Bank (funded by the Ontario Institute for Cancer Research), the FRQS tissue bank (Université de Sherbrooke), the Alberta Research Tumour Bank. Patient information and consent were managed by the tumour banks and released for this study under their respective material transfer agreements. Tissue handling and experimental procedures were approved by the Comité d’éthique de la recherche du CIUSSS de l’Estrie–CHUS (CÉR #10-062, 2010-241). Sample provenance, characteristics, RNA extraction, integrity and quality controls were described previously [16]. The discovery-cohort tissues from that study were re-sequenced here by poly(A)-selected RNA sequencing.

### RNA sequencing

Poly(A)-selected libraries were prepared from 500 ng total RNA from the tissue samples and the CRISPR-Cas13d-mediated SNORA81 depletion experiments using the NEBNext Ultra II Directional RNA Library Prep Kit (E7760, New England Biolabs) with 10–12 amplification cycles. Libraries from the tissue samples were sequenced on an Illumina instrument using a High Output 75-cycle cartridge with single-end 1 × 80 bp. Libraries from CRISPR–Cas13d-mediated SNORA81 depletion were sequenced on an Element Biosciences AVITI system using the Cloudbreak Freestyle High Output kit (paired-end 2 × 75 bp). TGIRT-seq libraries from ASO-mediated SNORA81 depletion were prepared as previously described [17–19]. RNA was ribodepleted with the Ribo-Zero Gold rRNA Removal Kit, purified using the Zymo RNA Clean & Concentrator with 8 volumes of ethanol to retain small RNAs, and used unfragmented for template-switching reactions with 1 µM TGIRT-III (InGex) for 15 min at 60 °C. cDNA was purified with the MinElute kit (QIAGEN), ligated at the 3′ end to a 5′-adenylated DNA oligonucleotide and amplified for 12 PCR cycles. Libraries were size-selected with AMPure XP beads (Beckman Coulter), assessed on a 2100 Bioanalyzer and sequenced on the Illumina system using a High Output 75-cycle cartridge with single-end 1 × 80 bp. ERCC RNA Spike-In Mix 1 (Ambion) was added to each sample as a control for detection uniformity.

### RNA sequencing analysis

All datasets were processed with a common pipeline [16]. Read quality was assessed with FastQC (Babraham Bioinformatics), and adapters and low-quality bases were removed with Trimmomatic [20]. Reads were aligned to hg38 with STAR v2.7.11b (--outFilterScoreMinOverLread 0.3, -- outFilterMatchNminOverLread 0.3, --outFilterMultimapNmax 100, --winAnchorMultimapNmax 100, --alignEndsProtrude 5 ConcordantPair) [21] using an index built from Ensembl v87 annotation supplemented with missing tRNA and snoRNA entries. Primary alignments were retained with SAMtools v1.22.1 [22]. Transcripts were quantified with CoCo correct_count v1.0.0, and differential expression was determined with DESeq2 using raw CoCo counts [23]. Transcripts with |log2 fold change| ≥ 0.5 and Benjamini–Hochberg-adjusted *P* ≤ 0.05 were considered differentially expressed. Volcano plots were generated with EnhancedVolcano (Bioconductor). Sample quality was evaluated using read quality, mapping rate, library complexity, inter-sample correlation and principal-component analysis. One tissue library (HGSC_3) and the libraries from one ASO replicate and one Cas13d replicate (g81_N2 and gSCR_N2) were excluded as global outliers, reflecting experiment-wide technical failure rather than biological variation. All exclusions were applied before differential-expression and enrichment analyses, and the number of samples retained per group is indicated in the figure legends.

### Cell culture and transfection

OVCAR-3 cells (CLS #300307/p690_NIH:Ovcar-3; RRID: CVCL_0465) were grown in RPMI 1640 supplemented with 20% (v/v) FBS and 2 mM L-glutamine at 37 °C under 5% CO₂, passaged according to ATCC recommendations for no more than 15 passages, and tested periodically for mycoplasma by qPCR (RNomics platform, Université de Sherbrooke) [16]. Cells were seeded at 1.0 × 10⁴ per well (96-well) or 3.5 × 10⁵ per well (six-well) and transfected with Lipofectamine 2000 (Invitrogen) according to the manufacturer’s instructions. Guide, ASO and plasmid sequences are listed in **Supplementary Table 1**. For SNORA81 depletion, cells received Cas13d and a pool of three SNORA81-targeting guide plasmids at a 1:1 mass ratio (1 000 ng total per six-well; 100 ng per 96-well), with a scrambled guide as control. For combined depletion, SNORA81-targeting guides were co-transfected with three IRAK1- or SPC24-targeting guides (600 ng per six-well; 60 ng per 96-well). ASO-mediated depletion used 15 nM SNORA81-targeting or control ASO composed of five 2′-O-methyl RNA bases, ten phosphorothioate DNA bases and five 2′-O-methyl RNA bases [16]. Cells were analysed at least 48 h after transfection.

### ITGB4 overexpression and rescue

ITGB4 was expressed from pRK5 β4 (Addgene 16037, deposited by F. G. Giancotti; [24]). Cells received 2 500 ng plasmid per six-well or 250 ng per xCELLigence well. For rescue experiments, cells were co-transfected with Cas13d, pooled SNORA81-targeting or scrambled guides, and either pRK5 β4 or empty pRK5 (3 500 ng total per six-well; 350 ng per xCELLigence well), with total DNA equalized using empty vector. The four conditions were gSCR plus empty vector, gSNORA81 plus empty vector, gSCR plus ITGB4 and gSNORA81 plus ITGB4. Cells were analysed at least 48 h after transfection.

### RNA extraction and RT-qPCR

RNA was extracted with the RNeasy Mini Kit (QIAGEN) as recommended, except that RNA was precipitated with 1.5 volumes of 100% ethanol to retain small RNAs, and assessed on a 2100 Bioanalyzer [16]. cDNA was synthesized from 500 ng–1 µg RNA using reverse transcriptase MMuLV (#MRT, Plateforme de purification des protéines de l’Université de Sherbrooke), RNase Inhibitor (20 units, #RI-L, Plateforme de purification des protéines de l’Université de Sherbrooke), dNTPs (1 mM) and random hexamers (3 µM). Reactions (10 µl) contained 5uLSupermix 2X (#S-mix) (Plateforme de purification des protéines de l’Université de Sherbrooke), 3 µl cDNA (diluted to 3,33ng/uL) and 200 nM of each primer, and were run in 384-well plates on a Bio-Rad CFX Real-Time System with no-template and no-reverse-transcriptase controls. Normalized relative expression was calculated by the ΔΔCq method against *MRPL19*, *RPPH1* and *YWHAZ*. Primers are listed in **Supplementary Table 1**.

### Western Blot

Cells were collected 48 h and 72 h after transfection and lysed in 50 µl buffer for 1 h at 4 °C with rotation, followed by centrifugation for 20 min, as previously described [16]. Protein was quantified by Bradford assay, and 30 µg was solubilized in Laemmli buffer (62.5 mM Tris-HCl pH 6.8, 10% glycerol, 5% β-mercaptoethanol, 2.3% SDS), denatured 5 min at 95 °C and resolved on 10–12% gels. Proteins were transferred to Amersham Protran nitrocellulose (Cytiva) for 90 min at 100 V, and membranes were blocked overnight at 4 °C in TBS containing 0.1% Tween-20 and 5% skimmed milk. Membranes were incubated for 90 min at room temperature with antibodies against Akt (1:2 000; 9272, Cell Signaling Technology), phospho-Akt Thr308 (1:1 000; 4056, Cell Signaling Technology), IRAK1 (1:2 000; 4504, Cell Signaling Technology), integrin β4 (1:500; 14803, Cell Signaling Technology), RASGRP2 (1:1 000; 30189-1-AP, Proteintech), SPC24 (1:5 000; ab169786, Abcam) and GAPDH (1:5 000; NB300-221, Bio-Techne), washed in TBS containing 0.1% Tween-20, and incubated with HRP-conjugated anti-rabbit (1:2,000; NA934, Cytiva) or anti-mouse (1:2 000; NA931V, Cytiva) secondary antibodies. Signals were detected with Clarity Western ECL (Bio-Rad) on a LAS-4000 imager (GE Healthcare). Band intensities were quantified by densitometry, normalized to GAPDH and expressed relative to the scrambled-guide control.

### Real-time cell growth

Cell adherence and growth were monitored using xCELLigence (ACEA Biosciences) E-plates [16]. Complete medium (50 µl) was added per well and plates were equilibrated for 30 min at 37°C under 5% CO₂ before addition of 1.0 × 10⁴ cells in 50 µl. Cell index was recorded every 10 min for 24 h to 48 h before transfection, until cells reached 70% confluence or a cell index of 0.6–1.0. Cells were transfected in-plate, cell index was normalized at the time of transfection, and recording continued every 10 min for at least 80 h and up to 120 h. Each experiment comprised three biologically independent replicates with three technical replicates.

### Cell-cycle analysis

Forty-eight hours after transfection, cells were washed in PBS, trypsinized, pelleted for 5 min at 100 g and resuspended in ice-cold PBS [16]. Cells were fixed by addition of ice-cold 100% ethanol and held at −20 °C for at least 30 min, then pelleted for 5 min at 400 g followed by 30 s at 16 200 g and washed in PBS, repeating the centrifugation and wash three times. Cells were treated with RNase A (100 µg/ml in PBS) for 5 min at room temperature and stained with propidium iodide for 30 min at room temperature. DNA content was measured on a BD LSRFortessa flow cytometer, and G1/G0, S and G2/M fractions were determined from DNA-content histograms. Experiments were performed in three biologically independent replicates.

### Wound-closure assay

Assays were performed 48 h after transfection in 96-well plates [16]. Medium was removed and a scratch was made across the centre of each well with a 200-µl pipette tip. Medium containing 60 mM hydroxyurea was then added to inhibit proliferation, so that wound closure reflected cell migration. Images were acquired with a CloneSelect imager (Molecular Devices) at 0, 24, 48 and 96 h. Wound closure was expressed as the percentage of the initial wound area covered. Experiments were performed in three biologically independent replicates.

### Pathway enrichment analysis

Over-representation analysis was performed with ShinyGO v0.85.2 [25] using a false-discovery-rate cutoff of 0.05, gene-set sizes between 2 and 5 000 genes, and display of the 20 most enriched terms. Gene sets were obtained from the KEGG database [26] and the MSigDB Hallmark collection [27]. KEGG pathway maps were rendered with Pathview [28].

### Statistical analysis

The number of biologically independent experiments is stated in each figure legend. Data are presented as mean ± s.d. Two-group comparisons used two-tailed Student’s *t*-tests assuming unequal variance [16]. Sequencing and enrichment *P* values were adjusted using the Benjamini– Hochberg procedure. *P* < 0.05 was considered significant.

## RESULTS

### SNORA81 depletion remodels PI3K/AKT-associated gene expression in HGSC

To identify the gene expression programmes through which SNORA81 acts, we compared HGSC and SBT tumour transcriptomes with two independent SNORA81-depletion datasets in OVCAR-3 cells, one generated using ASOs and profiled by TGIRT-seq following ribodepletion and the other generated using CRISPR/Cas13d and profiled by poly(A) RNA sequencing (**Figure 1B**). Requiring |log2 fold change| ≥ 0.5 and an adjusted *P* ≤ 0.05, 2 179 mRNAs differed between HGSC and SBT tissues, of which 910 were more abundant and 1 269 less abundant in HGSC. SNORA81 depletion altered 350 mRNAs by ASO and 117 mRNAs by CRISPR/Cas13d (**Figure 1B, C**). Over-representation analysis of these datasets against the MSigDB Hallmark collection returned interferon alpha response, E2F targets and TNFA signalling via NF-κB as the terms with the highest fold enrichment, together with the p53 pathway, interferon gamma response and PI3K/AKT/mTOR signalling (FDR < 0.05; **Supplementary Figure 1A**). Each depletion approach resulted in both increased and decreased mRNA abundance, indicating remodelling of specific gene expression programmes rather than a general loss of RNA.

Comparison of the three datasets identified 55 mRNAs differentially expressed in the tissue comparison and after at least one method of SNORA81 depletion, of which 35 were shared with the ASO dataset only, 15 with the CRISPR/Cas13d dataset only and 5 with both (**Figure 1C and Supplementary Figure 1B**). Twenty-two of the 55 mRNAs showed reversal of the HGSC-associated pattern after depletion of SNORA81, with transcripts more abundant in HGSC reduced after depletion and transcripts less abundant in HGSC increased (**Figure 1C and Supplementary Figure 1C**). For this analysis, the tissue comparison was oriented as SBT versus HGSC rather than HGSC vs SBT, so that the direction of the log2 fold change was aligned with the direction of the ASO- and CRISPR/Cas13d-depletion datasets. Thus, mRNAs more abundant in HGSC and reduced following SNORA81 depletion, or less abundant in HGSC and increased following depletion, have matching log2 fold-change signs across datasets (**Supplementary Figure 1C)**. Curation against KEGG PI3K–AKT annotation (hsa04151) linked 19 of the 22 mRNAs to this pathway; the remaining three (*VCP*, *MYO9A* and *WHRN*) are not represented in the map (**Figure 1C and Supplementary Figure 2A**).

To test whether these RNA changes reflected altered pathway activity, we measured AKT phosphorylation at Thr308 after CRISPR/Cas13d-mediated SNORA81 depletion. Depletion reduced the ratio of phosphorylated to total AKT to 0.30 of the scrambled guide control (*P* = 0.02, n = 3 biologically independent experiments), reflecting lower phosphorylated AKT (0.53 ± 0.06) and higher total AKT (1.89 ± 0.07) (**Figure 1D**). Reduced AKT phosphorylation in the absence of any loss of AKT protein identifies PI3K/AKT signalling as a pathway output of SNORA81 loss and places the primary regulatory event upstream of AKT.

### IRAK1 and SPC24 are downstream effectors of the SNORA81-dependent pathway state

We next asked whether these transcripts contribute to the invasive phenotype of SNORA81-depleted cells or merely report it. We therefore examined the 22 mRNAs whose HGSC-associated direction was reversed by SNORA81 depletion. Nineteen mapped to the KEGG PI3K–AKT pathway, distributed across NF-κB/stress crosstalk, cell-cycle and survival branches, with the direction of RNA change after SNORA81 depletion annotated on the map (**Supplementary Figure 2A**). We validated by RT-qPCR the RNA abundance of the 22 SNORA81-dependent mRNAs following SNORA81 depletion using the CRISPR/Cas13d system (**Supplementary Figure 2B**). Four candidates showed significant changes: *IRAK1* (*P* = 0.03), *SPC24* (*P* = 0.002) and *PIK3C2B* (*P* = 0.002) were significantly downregulated following SNORA81 knockdown, while *SOCS1* (*P* = 0.03) was the only candidate significantly upregulated.

We selected *IRAK1* and *SPC24* for functional validation because they showed the largest changes in mRNA abundance among the candidates selected for validation. *IRAK1* is a component of signalling downstream of the interleukin 1 receptor and has been implicated in immune response and inflammation by acting through Toll-like receptor-mediated signalling pathways and regulating interleukin (IL)-1R signalling [29]. *IRAK1* is also highly expressed in epithelial ovarian cancer (EOC) and has been implicated in HGSC proliferation, stemness and tumour growth [30]. *SPC24* is a component of the NDC80 kinetochore complex and is required for the accurate segregation of chromosomes during mitosis by coupling the kinetochore to spindle microtubules [31]. In cancer, *SPC24* has been implicated in breast cancer progression through PI3K/Akt signalling, and its knockdown increases apoptosis and attenuates cell growth and cell cycle progression [32]. These established functions provide a rationale to test whether downregulation of *IRAK1* and *SPC24* could phenocopy cellular effects observed following SNORA81 depletion. *PIK3C2B*, a node within the PI3K signalling pathway, is also significantly decreased following SNORA81 depletion, further supporting the dysregulation of this pathway.

We first assessed whether the change in *IRAK1* and *SPC24* following SNORA81 depletion extended to the protein level. Consistent with the RT-qPCR results, SNORA81 depletion, using Cas13d guides (gSNORA81), reduced both *IRAK1* and *SPC24* protein levels, to 0.62 ± 0.07 and 0.52 ± 0.09 of control, respectively (**Supplementary Figure 2C**). We next examined the specificity of targeting IRAK1 and SPC24 using Cas13d guides (gIRAK1 and gSPC24) in OVCAR-3 cells, relative to the scrambled guide control (gSCR). SNORA81 depletion reduced SNORA81 RNA abundance to 0.61 of control and lowered *IRAK1* to 0.63 and *SPC24* to 0.83, whereas depletion with gIRAK1 or gSPC24 reduced the abundance of their respective targets to 0.50 and 0.25, without affecting SNORA81 RNA levels (1.08 and 0.98) (**Figure 2A**). We then combined gSNORA81 with either gIRAK1 or gSPC24 to test whether their effects were additive. The combination of target-specific guides produced similar knockdown effects to the individual guides, despite each guide being delivered at half the single-guide input, SNORA81 was reduced to 0.54 and 0.52 and *IRAK1* to 0.49, while *SPC24* depletion was weaker in combination than alone (0.52 versus 0.25) (**Figure 2A**).

**Figure 2.**
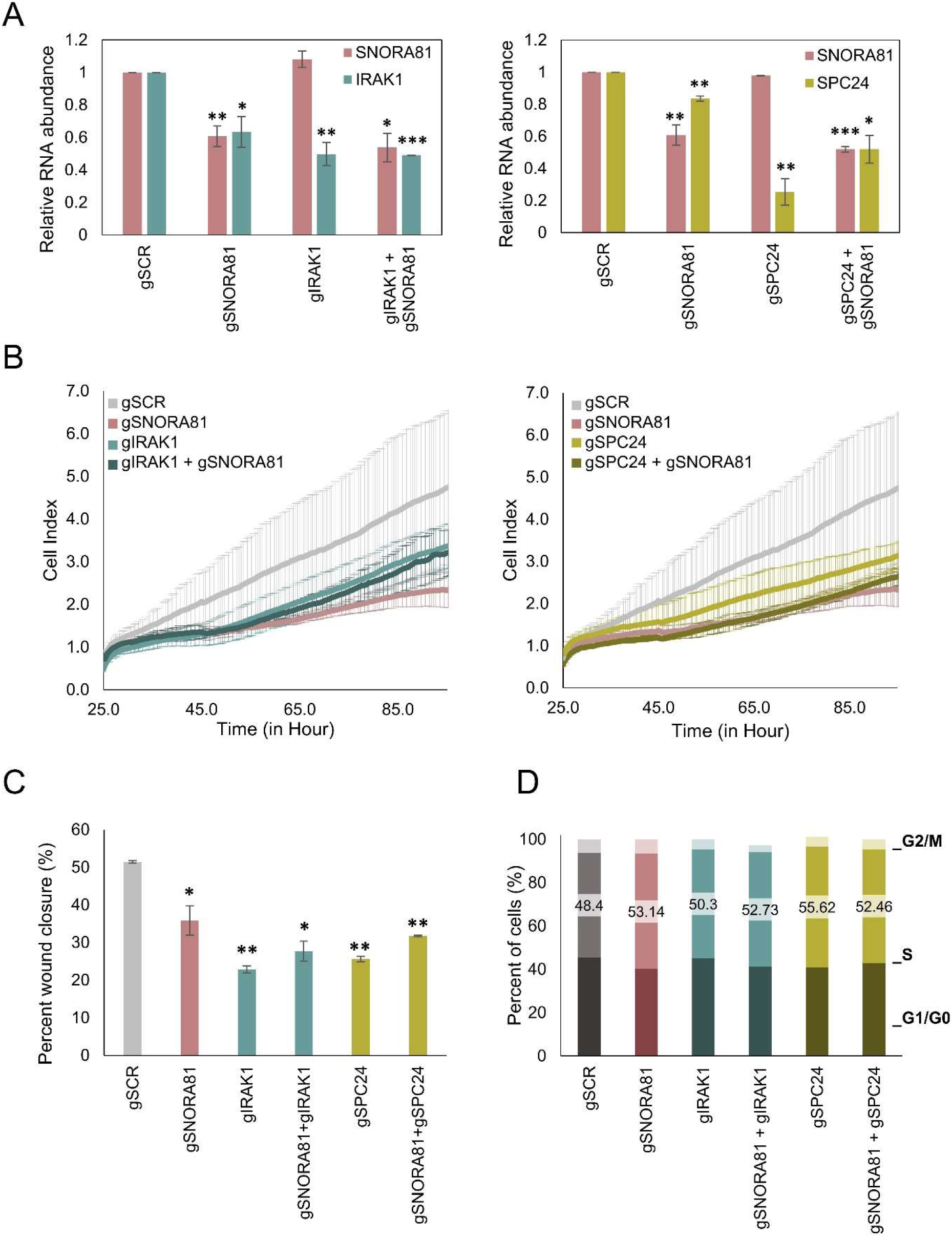
IRAK1 and SPC24 are downstream effectors of the SNORA81-dependent pathway state. All experiments used OVCAR-3 cells transfected with Cas13d and the indicated guide RNAs. gSCR, grey; gSNORA81, pink; gIRAK1, teal; gSPC24, olive;gSNORA81/gIRAK1, darker teal; gSNORA81/gSPC24, darker olive. **A**, SNORA81 depletion reduces *IRAK1* and *SPC24* RNA abundance, whereas depletion of *IRAK1* or *SPC24* does not affect SNORA81. Relative RNA abundance measured by RT-qPCR 48 h after transfection, normalised to *MRPL19*, *RPPH1* and *YWHAZ* and expressed relative to gSCR set to 1. Mean ± s.d.; n = 3 biologically independent experiments; two-tailed unpaired *t*-test against gSCR; \**P* < 0.05, \*\**P* < 0.01, \*\*\**P* < 0.001. **B**, SNORA81 depletion reduces cell growth more than depletion of *IRAK1* or *SPC24*, and combined depletion does not further reduce cell growth relative to either single depletion. Real-time cell index measured by xCELLigence and normalised at transfection time point; time is shown from seeding, with transfection at 24 h. Curves show the mean of n = 3 technical replicates. **C**, Depletion of SNORA81, *IRAK1* or *SPC24* reduces wound closure. Wound healing assay summarized as percent wound closure after 96 h in the presence of 60 mM hydroxyurea; n = 3 biologically independent experiments. **D**, SNORA81 and SPC24 depletion increase the proportion of cells in S phase. Cell-cycle distribution determined by propidium iodide staining and flow cytometry 48 h after transfection, showing the percentage of cells in G1/G0, S and G2/M.

We then assessed the impact of individual and combined knockdown on cell growth using real-time cell index measurements (xCELLigence). Depletion of either *IRAK1* or **SPC24**, as compared with control gSCR, reduced cell growth, but less than the effect of SNORA81 depletion (**Figure 2B**). Combination of gSNORA81/gIRAK1 and gSNORA81/gSPC24 had a similar impact to the individual guides, suggesting an epistatic relationship between downstream RNA-response genes in cell growth. We then examined cell migration using a wound healing assay performed in the presence of hydroxyurea to inhibit proliferation, observing cell migration for up to 96 hours. In our control, gSCR, cells progressively covered the wound, reaching 51.4% closure at 96 h, whereas SNORA81, *IRAK1* or *SPC24* depletion reduced wound coverage to 35.9%, 22.9% and 25.6% respectively (**Figure 2C**). Combined gSNORA81/gIRAK1 and gSNORA81/gSPC24 depletion had similar effects on wound closure to those observed with depletion of each target individually (27.7% and 31.7%). SNORA81 depletion therefore masks the contribution of each effector. Finally, we measured cell cycle progression following depletion of SNORA81, *IRAK1* or *SPC24*. In both arms, S-phase fraction was 48.4% for gSCR and 53.1% for gSNORA81. In the IRAK1 arm, S-phase fractions were 50.3% for gIRAK1 and 52.7% for gSNORA81/gIRAK1. In the SPC24 arm, S-phase fractions were 55.6% for gSPC24 and 52.5% for gSNORA81/gSPC24. SNORA81 and *SPC24* depletion increased the proportion of cells in S phase relative to the control, but *IRAK1* depletion alone did not produce this effect (**Figure 2D**). Although cell-cycle arrest has been reported following *IRAK1* depletion, the effect of *IRAK1* on migration and proliferation may occur through mechanisms that do not result in a large steady-state shift in cell-cycle distribution in OVCAR-3 cells [33]. The absence of additive effects on growth, migration and cell-cycle distribution places *IRAK1* and *SPC24* downstream of SNORA81 rather than in parallel with it, identifying them as effectors of the SNORA81-dependent pathway response rather than its cause.

### SNORA81 depletion reduces translation of upstream PI3K/AKT activators

SNORA81 guides an rRNA modification and is not predicted to interact directly with the mRNAs whose abundance changes upon its depletion, including *IRAK1* and *SPC24*. We therefore hypothesised that the PI3K/AKT changes described above are triggered by SNORA81-dependent changes in translation, acting on components upstream of AKT. To test this, we examined the translational regulation of upstream PI3K/AKT pathway components after SNORA81 depletion. Because changes in RNA abundance may represent secondary responses to altered signalling, we focused on SNORA81-dependent translationally altered RNAs (STAR mRNAs), defined by altered polysome association after CRISPR/Cas13d-mediated SNORA81 depletion under conditions in which ribosome integrity and global translation are unaffected [34]. Among 13 025 input-expressed mRNAs analysed by polysome profiling, 524 STAR mRNAs were identified, comprising 289 under-associated and 235 over-associated transcripts (**Figure 3A**). These changes were largely independent of mRNA abundance, with 485 STAR mRNAs remaining after filtering against matched RNA-seq data. Of these 485 transcripts, 29 corresponded to PI3K/AKT pathway components, including genes involved in growth factor, integrin/adhesion, RAS/MAPK and receptor/cytokine signalling (**Figure 3A**). We further prioritised 17 candidates positioned upstream of AKT activation. All 17 were under-associated, a direction consistent with the reduced AKT phosphorylation observed following SNORA81 depletion, and included growth factors and regulators of signalling, cell motility and growth (**Supplementary Figure 3A**).

**Figure 3.**
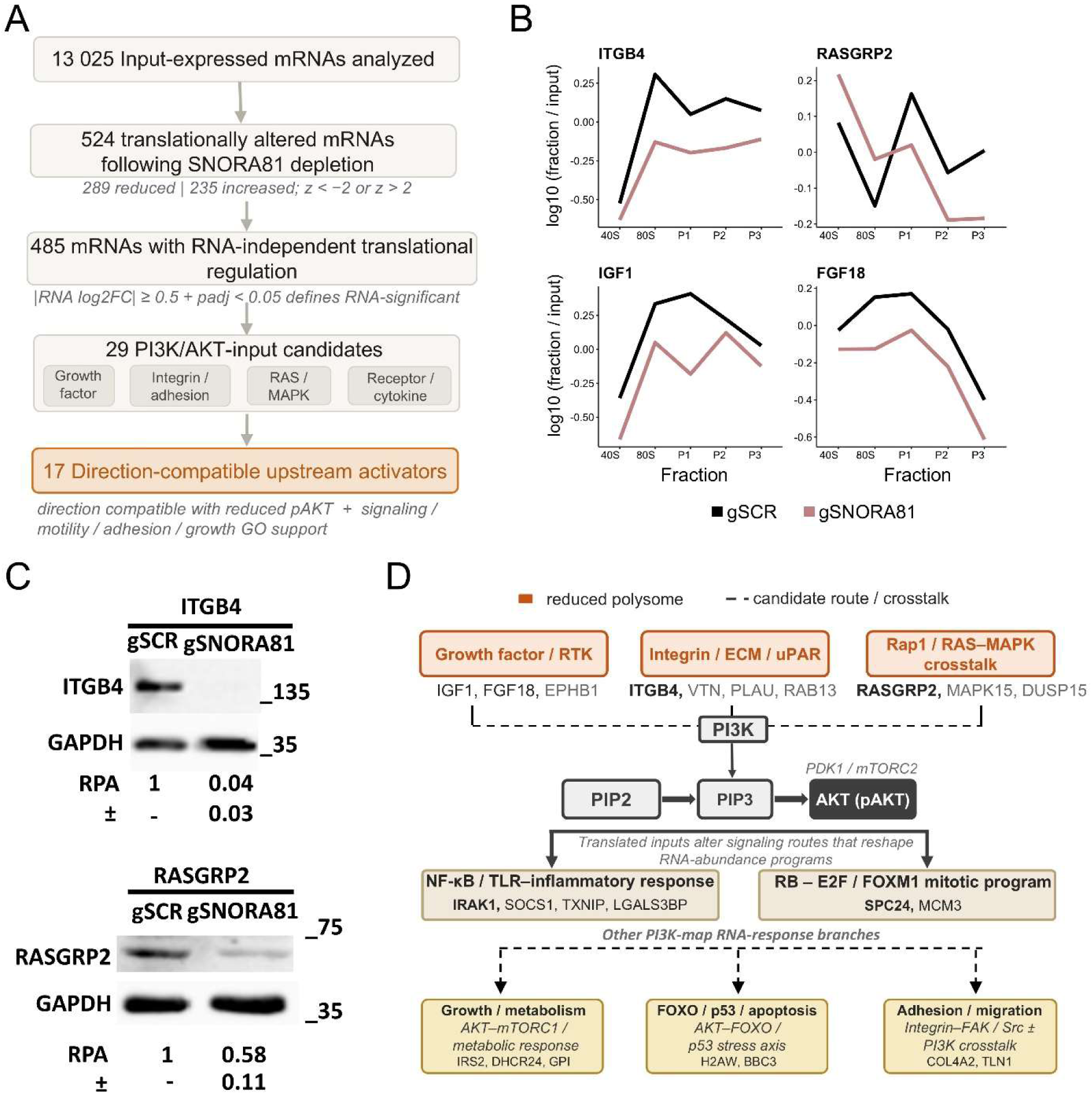
SNORA81 depletion reduces translation of upstream PI3K/AKT activators. **A**, Workflow for identifying SNORA81-dependent translationally altered candidates. Of 13 025 input-expressed mRNAs, 524 showed altered polysome association after SNORA81 depletion (289 reduced, 235 increased). Filtering against matched RNA-abundance data identified 485 transcripts with RNA-independent translational regulation, of which 29 corresponded to PI3K/AKT pathway components. Seventeen upstream activators with reduced polysome association were prioritised as direction-compatible with the reduced AKT phosphorylation in Fig. 1D. Selection thresholds are indicated in the panel. Polysome profiling data are from Faucher-Giguère *et al.* (submitted) [34]. **B**, SNORA81 depletion reduces ribosome association of *ITGB4*, *RASGRP2*, *IGF1* and *FGF18*. Distribution of each mRNA across sucrose-gradient fractions in gSCR control (black) and SNORA81-depleted (pink) cells, expressed as log10 fraction TPM relative to the matched input sample. P1, two to three ribosomes; P2, four to six; P3, seven or more. n = 3 biologically independent experiments. **C**, SNORA81 depletion reduces ITGB4 and RASGRP2 protein abundance. Western blots probed for ITGB4 or RASGRP2, with GAPDH as loading control. Relative protein abundance was normalised to GAPDH and expressed relative to gSCR: ITGB4, 0.04 ± 0.03; RASGRP2, 0.58 ± 0.11. Mean ± s.d.; n = 3 biologically independent experiments; two-tailed unpaired *t*-test, ITGB4 *P* = 0.0003, RASGRP2 *P* = 0.02. Molecular weights are indicated in kDa, and blots were cropped for clarity. **D**, Model integrating translational regulation of upstream activators with downstream RNA-abundance programmes. Activators are grouped by pathway entry point. Orange, reduced polysome association after SNORA81 depletion; bold, candidates validated at the protein level in this study; plain type, additional candidates from the 29-candidate set; dashed lines, candidate routes and crosstalk. Downstream modules summarise the NF-κB/TLR-inflammatory response, the RB–E2F/FOXM1 mitotic programme, and growth/metabolism, FOXO/p53/apoptosis and adhesion/migration branches.

We selected four of these candidates for further analysis: insulin-like growth factor 1 (*IGF1*), integrin subunit beta 4 (*ITGB4*), fibroblast growth factor 18 (*FGF18*) and Ras guanyl releasing protein 2 (*RASGRP2*). These candidates lie upstream of AKT activation and are linked to the cellular processes altered by SNORA81 depletion, namely growth, adhesion and motility. Polysome profiles for all four showed reduced association with the heavy polysome fractions following SNORA81 depletion (**Figure 3B**). We next tested whether reduced ribosome association translated into reduced protein output. *ITGB4* is a receptor that contributes to cell adhesion, signalling, migration and invasion, and affects EOC progression in OVCAR-3 cells [35, 36], whereas *RASGRP2* activates Ras and has been implicated in pathways relevant to cancer progression [37, 38]. SNORA81 depletion reduced ITGB4 and RASGRP2 protein abundance to 0.04 ± 0.03 (*P* = 0.0003) and 0.58 ± 0.11 (*P* = 0.02) of control, respectively (**Figure 3C and Supplementary Figure 3B**). IGF1 and FGF18 therefore remain polysome-supported candidates, whereas ITGB4 and RASGRP2 provide protein-level confirmation that SNORA81 loss reduces the output of upstream pathway inputs.

Together, these findings indicate that SNORA81 depletion reduces the translational engagement and protein output of upstream PI3K/AKT activators. These inputs comprise growth factor/RTK signalling, integrin/ECM/uPAR signalling, Rap1/RAS–MAPK crosstalk and additional context-dependent inputs, all converging on PI3K activation, PIP2-to-PIP3 conversion and AKT phosphorylation (**Figure 3D**). The same map places the resulting loss of pathway activity upstream of the NF-κB/TLR-inflammatory response and the RB–E2F/FOXM1 mitotic programme, which contain *IRAK1* and *SPC24* (**Figure 3D**). The two layers are affected at different levels: upstream activators lose ribosome association and protein output, whereas downstream effectors change only in RNA abundance. SNORA81-dependent rRNA modification therefore acts as a translational filter on PI3K/AKT pathway entry, coupling ribosome state to oncogenic signalling output.

### ITGB4 restoration partially suppresses the SNORA81-depletion phenotype

Loss-of-function experiments establish that SNORA81 is required for pathway activity but cannot place its point of action within the pathway. We therefore asked whether restoring a single translationally repressed activator downstream of the SNORA81 lesion suppresses the depletion phenotype, and tested this for ITGB4, the most strongly reduced activator at the protein level (**Figure 4A**). We first confirmed increased ITGB4 protein following transfection of the ITGB4 expression construct, which raised ITGB4 abundance 5.6-fold relative to empty vector (**Figure 4B**). We next evaluated cell growth and found that *ITGB4* overexpression partially recovered the growth defects associated with SNORA81 depletion, with cell index at 103 h recovering from 2.06 to 2.89 against 3.20 in cells receiving the scrambled guide and empty vector (**Figure 4C**). Finally, we assessed the expression of the downstream SNORA81-dependent response genes *IRAK1* and *SPC24* following ITGB4 overexpression. Their RNA abundance, which fell to 0.45 and 0.48 of control after SNORA81 depletion, was restored following ITGB4 overexpression in SNORA81-depleted cells, reaching 8.4 and 3.3 respectively (**Figure 4D**). Suppression of the SNORA81 phenotype by ITGB4 expression, which cannot restore the rRNA modification itself, places ITGB4 downstream of SNORA81 and upstream of the IRAK1 and SPC24 response. The partial recovery is consistent with pathway flux being controlled by several translationally repressed activators rather than vested in one.

**Figure 4.**
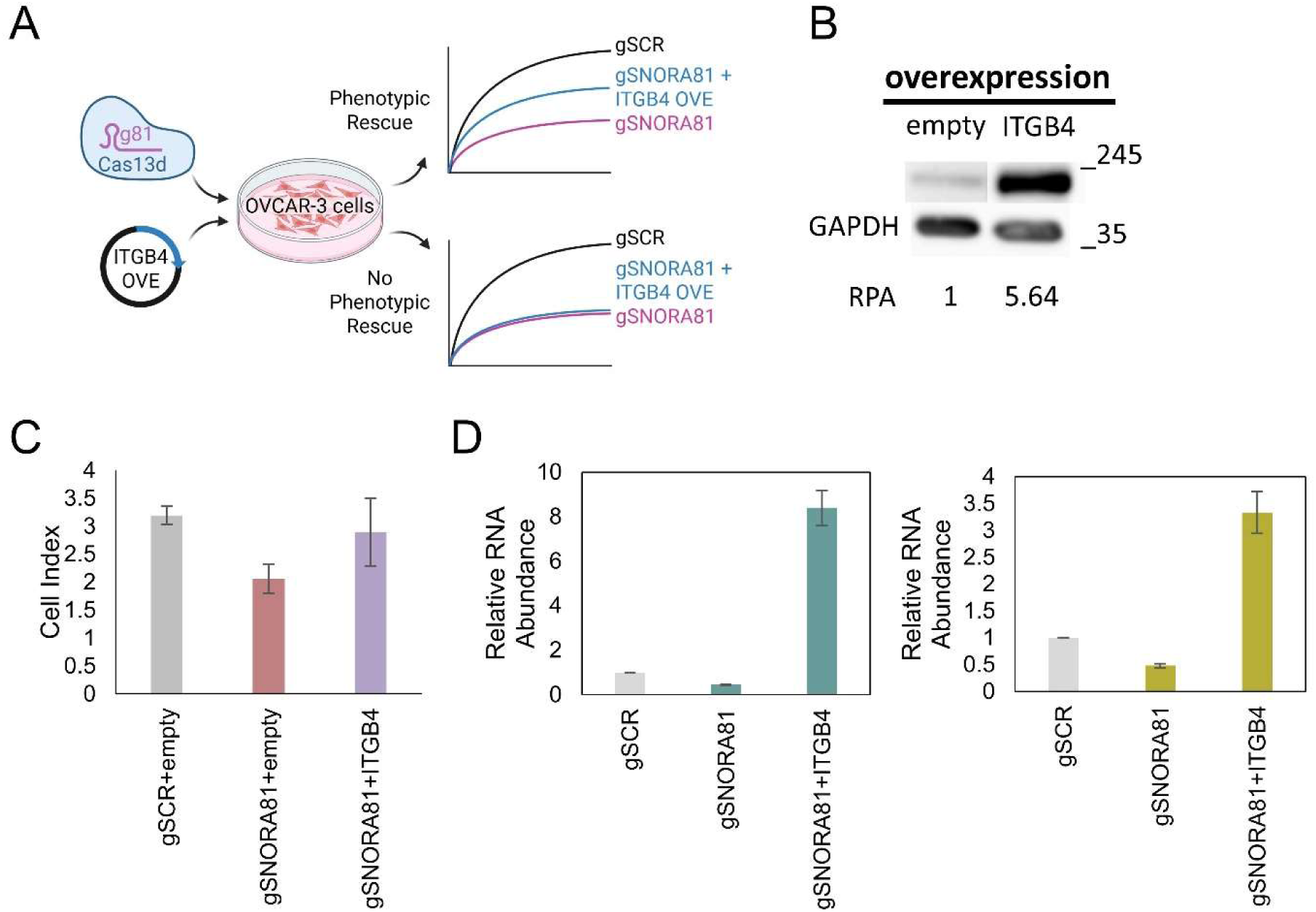
ITGB4 restoration partially suppresses the SNORA81-depletion phenotype. **A**, Experimental design. OVCAR-3 cells were co-transfected with Cas13d and either SNORA81-targeting guides (gSNORA81) or a scrambled control guide (gSCR), together with an ITGB4 expression construct or empty pRK5 vector. **B**, Validation of ITGB4 overexpression. Western blot of ITGB4 in cells transfected with empty pRK5 or 2.5 µg ITGB4 expression construct, with GAPDH as loading control; relative protein abundance is shown beneath the blot. **C**, ITGB4 expression partially restores cell growth after SNORA81 depletion. Real-time cell index measured by xCELLigence in cells transfected with gSCR plus empty vector, gSNORA81 plus empty vector, or gSNORA81 plus ITGB4. Time is shown from seeding, with transfection at 24 h. Bars show mean ± s.d at 103 h. **D**, ITGB4 expression restores *IRAK1* and *SPC24* RNA abundance after SNORA81 depletion. Relative RNA abundance measured by RT-qPCR, normalised to *MRPL19*, *RPPH1* and *YWHAZ* and expressed relative to empty pRK5 vector.

### SNORA81-dependent rRNA modification establishes a translational layer of PI3K/AKT control in HGSC

Our data support the following model (**Figure 5**). In HGSC, increased SNORA81 raises 28S rRNA pseudouridylation level at U4636, generating a ribosome population that favours translation of the upstream PI3K/AKT activators *ITGB4*, *RASGRP2*, *IGF1* and *FGF18*. These activators enter the pathway through integrin/ECM, RAS–MAPK and growth factor/RTK routes and converge on PIP3 production and AKT phosphorylation. SNORA81 depletion reverses each step, reducing polysome association and protein abundance of these activators and lowering AKT phosphorylation without loss of AKT protein. Reduced pathway activity propagates to transcriptional response programmes, including NF-κB/TLR-inflammatory signalling containing *IRAK1* and the RB–E2F/FOXM1 mitotic programme containing *SPC24*, and to the growth, migration and cell-cycle phenotypes that characterise HGSC. Because the two layers act on separate gene sets and in a fixed order, a single rRNA modification can filter PI3K/AKT pathway entry at the level of translation.

**Figure 5.**
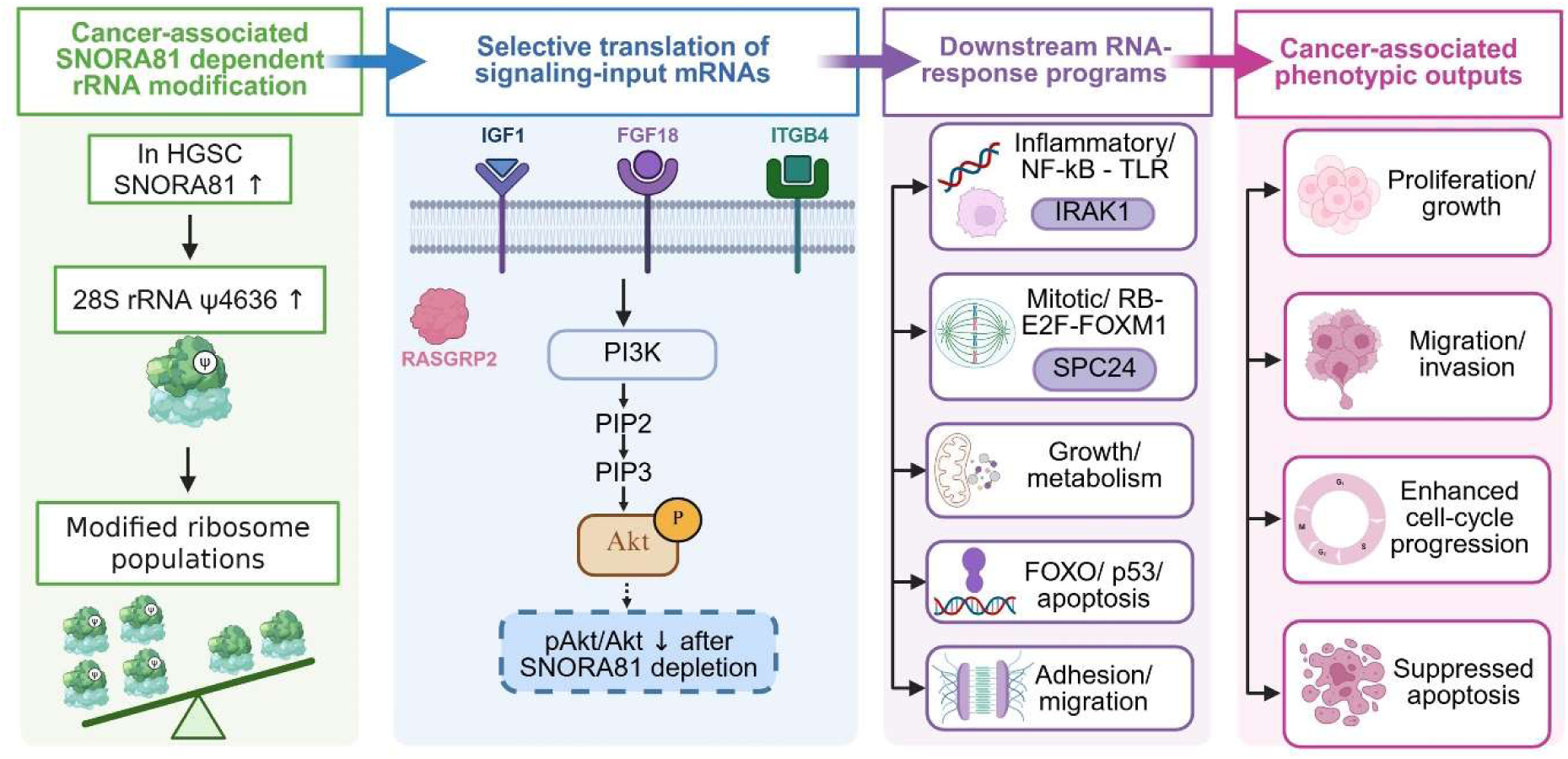
SNORA81-dependent rRNA modification establishes a translational layer of PI3K/AKT control in HGSC. Proposed model. In HGSC, increased SNORA81 raises pseudouridylation of 28S rRNA at U4636, generating a modified ribosome population that favours translation of mRNAs encoding upstream PI3K/AKT activators, including *IGF1*, *FGF18*, *ITGB4* and *RASGRP2*. These activators enter the pathway through growth factor/RTK, integrin/ECM and RAS–MAPK routes and converge on PIP3 production and AKT phosphorylation by PDK1 and mTORC2. SNORA81 depletion reduces their ribosome association and protein abundance and lowers the phospho-AKT/AKT ratio. Reduced pathway activity is proposed to remodel downstream RNA-response programmes, including NF-κB/TLR-inflammatory signalling containing *IRAK1*, the RB–E2F/FOXM1 mitotic programme containing *SPC24*, and growth/metabolism, FOXO/p53-dependent apoptosis and adhesion/migration branches. These programmes are associated with the proliferation, migration and invasion, and cell-cycle phenotypes of HGSC, and with the reduced apoptosis reported previously [16].

## DISCUSSION

We show that a snoRNA-guided rRNA modification regulates oncogenic signalling in HGSC by controlling the translation of a pathway’s upstream activators. Cancer signalling is usually attributed to genetic alteration, transcriptional change or post-translational control of signalling proteins [5]. Our data add a translational layer: SNORA81 loss reduces polysome association and protein output of upstream PI3K/AKT activators, lowers AKT phosphorylation, and remodels downstream transcriptional programmes without a general reduction in RNA abundance (**Figure 5**).

The architecture of this response has two properties that constrain its interpretation. First, the layers are separable. Upstream activators change at the level of ribosome association and protein output, whereas downstream effectors change only in RNA abundance, with corresponding changes in protein output. Selective translation of pathway inputs therefore precedes the broader RNA response rather than accompanying it. Second, the layers are ordered. ITGB4 expression cannot reconstitute the rRNA modification, yet it suppresses both the growth defect and the loss of *IRAK1* and *SPC24* expression, placing ITGB4 downstream of SNORA81 and upstream of the transcriptional response. Because PI3K/AKT signalling is among the most frequently dysregulated networks in ovarian cancer yet its activity is poorly predicted by canonical genomic alterations [5], a translational determinant of pathway input offers one explanation for that discrepancy.

Four activators entering through integrin/ECM, RAS–MAPK and growth factor/RTK routes showed reduced ribosome association after SNORA81 depletion, and ITGB4 and RASGRP2 also showed reduced protein abundance (**Figure 3C**). The involvement of multiple pathway entry points argues that SNORA81 does not act through a single signalling target. It also provides a rationale for why restoring ITGB4 alone suppresses only part of the phenotype: where control of pathway flux is shared among several convergent inputs rather than dominated by a single component. Restoring one input may leave the remaining repressed activators, limiting pathway output. This raises the possibility that modest translational changes distributed across convergent inputs may alter pathway output more effectively than a large change in any one of them. IGF1 and FGF18 remain candidates at this level; their reduced polysome association identifies them as translationally regulated pathway inputs, but protein-level validation is still required.

Downstream, *IRAK1* and *SPC24* behave as response genes rather than regulators. Depletion of either reduced growth and wound closure, and *SPC24* depletion also increased the S-phase fraction (**Figure 2**), consistent with the roles of *IRAK1* in inflammatory signalling and ovarian cancer growth [29, 30] and of *SPC24* in kinetochore function and PI3K/AKT-dependent proliferation [31, 32]. Their phenotypes were not identical, indicating that the downstream response is not a single uniform programme: *IRAK1* connects the SNORA81-dependent state to NF-κB/TLR signalling, whereas *SPC24* connects it to the mitotic programme (**Figure 3D**). Co-depletion of either gene with SNORA81 produced no additive effect, an alleviating genetic interaction that places both within the same pathway and downstream of SNORA81. That *IRAK1* depletion reduced migration without shifting cell-cycle distribution is consistent with reports that *IRAK1* acts on motility through mechanisms distinct from cell-cycle arrest [33].

These findings reframe rRNA modification as an active contributor to pathway regulation rather than a passive feature of ribosome biogenesis. Ribosomes are increasingly recognised as heterogeneous machines whose modifications may bias translation of particular mRNAs [12], and dysregulated snoRNA expression is a recurrent feature of tumours [15]; our data connect these observations by showing that a change at one rRNA site propagates through selected signalling inputs to reshape pathway output and cellular behaviour. Protein-level validation of IGF1 and FGF18, together with identification of the remaining translationally repressed activators, will determine how much of the SNORA81 phenotype is attributable to translational control of pathway entry. Whether such control extends beyond PI3K/AKT, and whether it can be exploited in tumours whose pathway activity is not predicted by genomic alteration, remain open. Because SNORA81 abundance distinguishes HGSC from serous borderline tumours and normal fallopian tube epithelium [16], translational control of pathway entry may prove to be a targetable feature of the aggressive disease state.

## ACKNOWLEDGEMENTS

The authors thank the members of the Scott and Abou Elela labs for insightful discussions.

## AUTHOR CONTRIBUTIONS

AR and LFG performed and analyzed the experiments. AR wrote the first draft of the manuscript, and LFG revised the manuscript. SAE and MSS conceived the study and contributed to the writing and revision of the manuscript.

## DATA AVAILABILITY

RNA sequencing data were deposited in the NCBI Gene Expression Omnibus (GEO; https://www.ncbi.nlm.nih.gov/geo) under the accession number GSE346589.

