## Supplementary Figures for "A Ribosomal RNA modification regulates translation of PI3K/AKT activators in high-grade serous ovarian carcinoma"

Rivera-Pizana et al. Supplementary figures

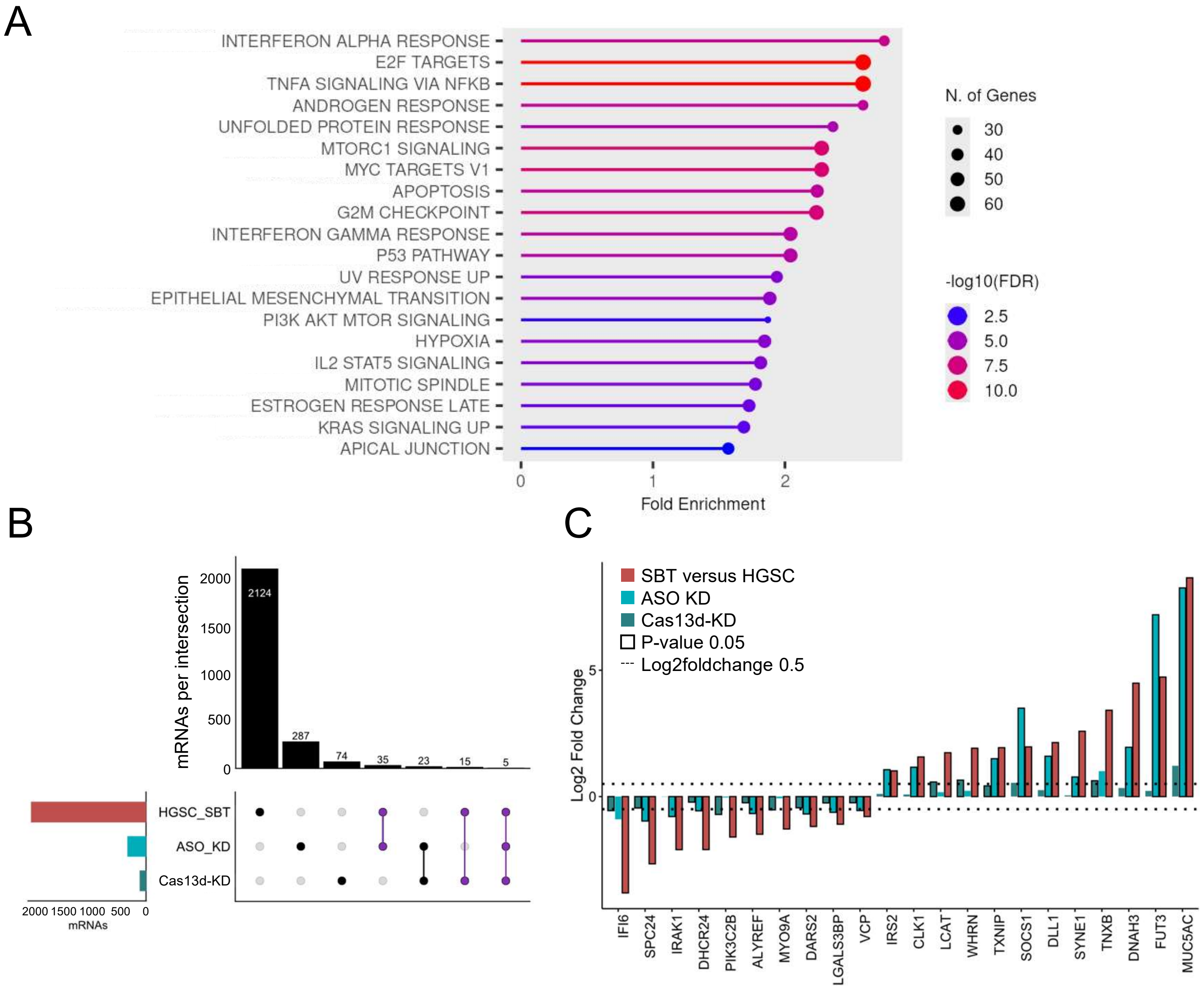


**Supplementary Figure 1. (Related to Figure 1) HGSC-associated pathways and mRNA overlap following SNORA81 depletion**. **A,** Lollipop plot of Hallmark analysis for the mRNAs differentially expressed in the SBT versus HGSC tissue comparison and in SNORA81-depletion datasets (ASO- or Cas13d-mediated knockdown). Over-representation testing was performed with ShinyGO v0.85.2 against the MSigDB Hallmark gene-set collection, using all expressed genes as the background, an FDR cutoff of 0.05 and gene-set sizes of 2–5 000 genes; the 20 most enriched terms are displayed. The x axis shows fold enrichment, dot size indicates the number of input genes assigned to each term, and colour indicates −log10(FDR). **B,** UpSet plot of differentially expressed mRNAs in SBT versus HGSC tissues, ASO-mediated SNORA81 knockdown, and CRISPR-Cas13d-mediated SNORA81 knockdown. Horizontal bars indicate the total number of differentially expressed mRNAs in each dataset, and vertical bars indicate the number of mRNAs in each intersection, defined by connected filled circles. Purple circles and connecting lines mark intersections containing HGSC/SBT mRNAs and at least one SNORA81-depletion dataset; these intersections comprise 55 shared mRNAs. **C,** Log2 fold changes for the 22 shared mRNAs showing opposite biological directionality between the HGSC/SBT comparison and SNORA81-depletion datasets. For this analysis, the tissue comparison was oriented as SBT versus HGSC, rather than HGSC versus SBT, so that its log2 fold-change direction was directly comparable with the ASO-KD and Cas13d-KD contrasts. Consequently, mRNAs elevated in HGSC and reduced following SNORA81 depletion, or reduced in HGSC and elevated following SNORA81 depletion, display concordant log2 fold-change signs across datasets. Bars are coloured by dataset. Black outlines indicate Benjamini-Hochberg-adjusted $P < 0.05$. For every mRNA, at least one SNORA81-depletion dataset was significantly differentially expressed, and the other depletion dataset showed the same direction of change. Dotted lines indicate absolute log2 fold change 0.5.

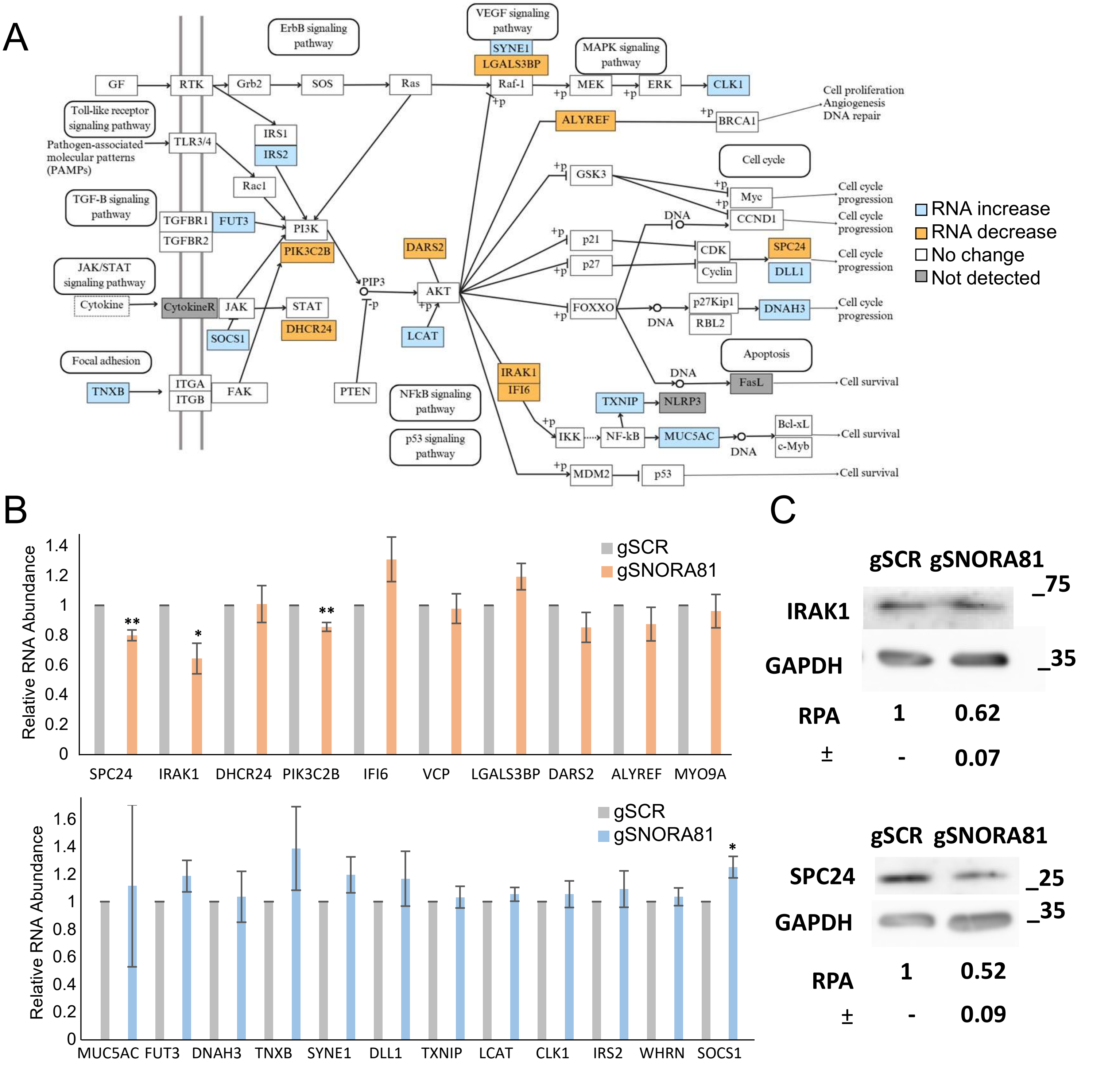


**Supplementary Figure 2. PI3K/Akt pathway assignment and validation of SNORA81-dependent RNA and protein changes (Related to Figure 1 and 2) A,** Modified KEGG PI3K/Akt signalling map (hsa04151) showing the 19 of 22 SNORA81-linked candidates that map to pathway components; *VCP*, *MYO9A* and *WHRN* are not represented. Blue indicates increased and orange indicates decreased RNA abundance after SNORA81 depletion, based on RNA sequencing. White indicates pathway components with no significant change, and grey indicates components not detected in the sequencing data. Candidates are positioned within receptor, integrin, PI3K, AKT, NF-κB/stress, apoptosis, cell-cycle and survival branches of the pathway. **B,** RT-qPCR validation of the 22 SNORA81-dependent mRNAs. Relative RNA abundance in gSCR and SNORA81-depleted OVCAR-3 cells, normalised to MRPL19, RPPH1 and YWHAZ and expressed relative to gSCR, set to 1. Candidates are grouped by the direction of change observed by RNA sequencing, with mRNAs predicted to decrease in the upper graph and those predicted to increase in the lower graph. Data are mean ± s.d. of three biologically independent experiments; two-tailed unpaired t-test against gSCR. Four candidates reached significance: *SPC24* (P = 0.002), *IRAK1* (P = 0.03), *PIK3C2B* (P = 0.002) and *SOCS1* (P = 0.03). **C,** Western blot validation of IRAK1 and SPC24 protein abundance following SNORA81 depletion. Blots were probed for IRAK1 or SPC24 and for GAPDH as a loading control. Relative protein abundance (RPA) was normalised to GAPDH and expressed relative to gSCR: IRAK1 0.62 ± 0.07 and SPC24 0.52 ± 0.09 (mean ± s.d., n = 3 biologically independent experiments). Molecular weight markers are indicated in kDa, and blots were cropped for clarity.

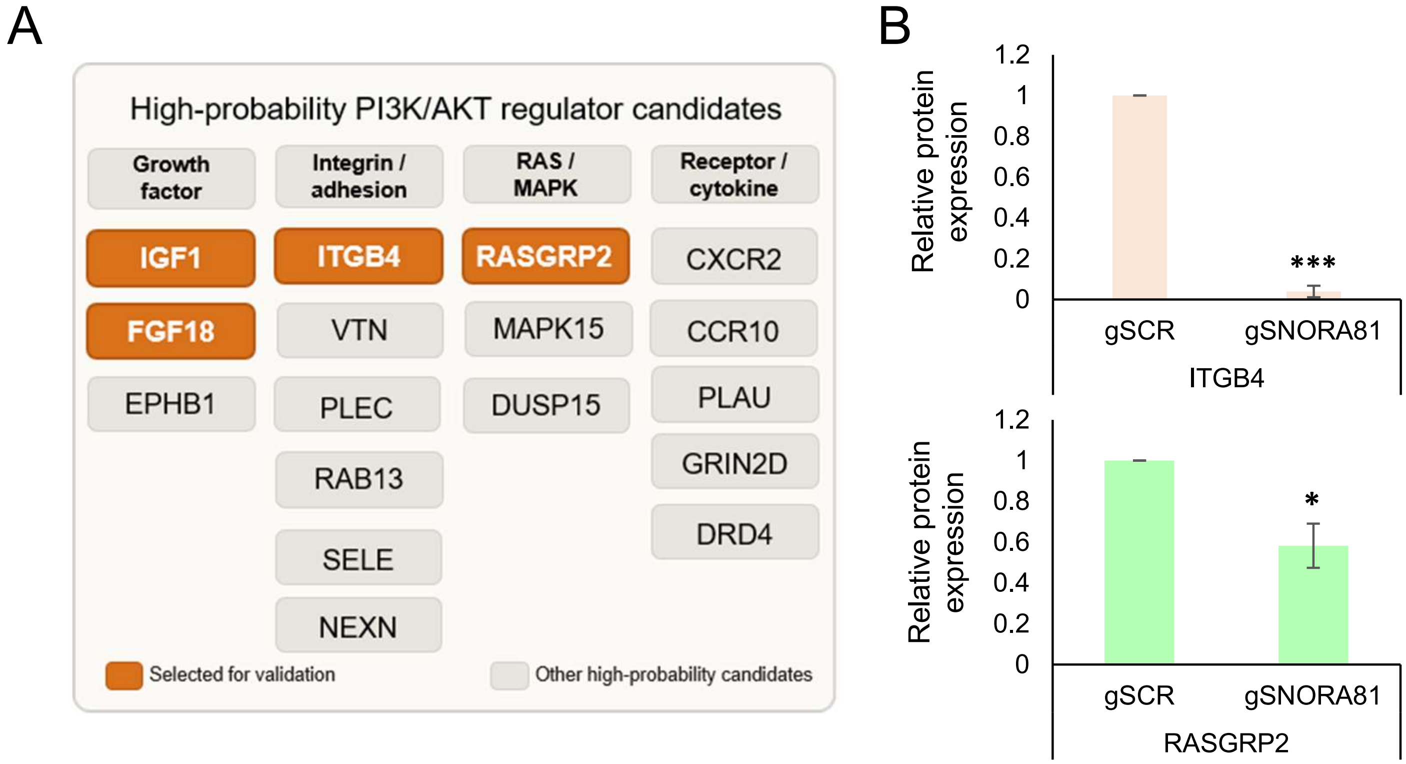


**Supplementary Figure 3. Classification of upstream PI3K/Akt candidates and quantification of ITGB4 and RASGRP2 protein abundance (Related to Figure 3) A,** The 17 direction-compatible upstream activators identified among SNORA81-dependent translationally altered (STAR) mRNAs, grouped by pathway entry point into growth factor, integrin/adhesion, RAS/MAPK and receptor/cytokine signalling. Orange indicates the four candidates selected from polysome profiling in Fig. 3B; grey indicates the remaining candidates. **B,** SNORA81 depletion reduces ITGB4 and RASGRP2 protein abundance. Quantification of the western blots shown in Fig. 3C for ITGB4 (top) and RASGRP2 (bottom) in gSCR control and SNORA81-depleted OVCAR-3 cells, normalised to GAPDH and expressed relative to gSCR. Data are mean ± s.d. of 3 biologically independent experiments; T-test against gSCR; ***P = 0.0003 for ITGB4, and *P = 0.02 for RASGRP2.
